# Patterns of secondary invasion in the understory of exotic, invasive timber stands

**DOI:** 10.1101/2022.11.29.518341

**Authors:** Varughese Jobin, Arundhati Das, C.P. Harikrishnan, Ritobroto Chanda, Swapna Lawrence, V.V. Robin

## Abstract

Current climate and land cover change threaten global mountaintops with increased spread of invasive species. Long-established plantations of exotic and invasive trees on these mountaintops can alter their surroundings, further increasing invader-facilitated or secondary invasion. Identifying the ecological conditions that promote such specific associations can help develop better management interventions.

The Western Ghats’s Shola Sky Islands (>1400m MSL) host vast stretches of exotic and invasive tree plantations that sustain colonisation of other invasive woody, herbaceous and fern species in their understories. Here we analysed vegetation and landscape variables from 232 systematically-placed plots in randomly selected grids using NMDS and Phi Coefficient approaches, to examine patterns of association (positive interactions) between secondary understory invasive species with specific exotic and invasive overstory species. We also conducted GLMM with zero inflation to determine the influence of environmental variables where such associations occur.

We find that secondary invasion of multiple species under the canopy of other exotic invasives is widespread across the Shola Sky Islands. Stands of Eucalyptus host the colonisation of 70% of non-native invasive species surveyed across the Shola Sky Islands. In particular, Lantana camara invasion is strongly associated with Eucalyptus stands.

We also found that climatic variables affect the colonisation of understorey woody invasive species, while invasion by exotic herbaceous species is associated with the density of road networks.. Canopy cover impacts all invasives negatively, while incidence of fire was negatively associated with invasion by *Lantana spp* and the *Pteridium spp*. While the restoration of natural habitats largely targets the highly invasive Acacia, less invasive Eucalyptus and Pine are often not included. Our study suggests that retaining such exotic species in natural habitats, particularly protected areas, can hinder ongoing restoration efforts by facilitating further invasions by multiple woody and herbaceous species.

## Introduction

Invasive plant species have great potential to expand within global biodiversity hotspots (J.-Z. Wan & Wang, 2018) and impact endangered species (Dueñas et al., 2021). Mountaintops face increasing threats from invasive species owing to the ongoing changes in climate and land cover (McDougall et al., 2011; Pauchard et al., 2009). Exotic invasive tree species have established extensive stands across several tropical mountaintops, or sky islands, during the past century (Arasumani et al., 2019; Hulme et al., 2013) posing serious threats to native ecosystems (Hejda et al., 2017). Apart from their negative impacts on native regeneration (Dyderski et al., 2017), studies from low-elevation ecosystems show that invasive tree stands can also display signs of heterospecific invasion (Kuebbing & Nuñez, 2015; Tecco et al., 2007). Studies on such invader-facilitated invasion or secondary invasion from mountains are few (Giantomasi et al., 2008) and, thus, warrant further exploration.

Some non-native trees that were introduced in grassland-forest mosaics in high-altitude mountains (Arasumani et al., 2019; Hulme et al., 2013) impact biodiversity, water and soil systems with their expansion into grasslands (Simberloff, 2011; van Wilgen & Richardson, 2014). Plantations of non-native tree species can alter their local habitat, and these effects get multiplied due to their size and longevity (Dyderski et al., 2017; Le Maitre et al., 2011). Their presence and influence may also facilitate the invasion of other non-natives (Kuebbing & Nuñez, 2015; O’Loughlin & Green, 2018). Such understory invasives can have cascading impacts by competing with natural forest regeneration (Dyderski et al., 2017; Vacek et al., 2020), impacting animal movements (Habel et al., 2016; Stewart et al., 2021) and enhancing edge effects (McDonald & Urban, 2006). Mapping such invasions at a landscape level becomes imperative to understand if particular overstory exotic and invasive species are associated with specific secondary invasive species in their understory. Large-scale management of such invasive tree stands on mountains can pose a great challenge to forest managers owing to topography and vegetation (McDougall et al., 2011), and identifying stands with significant understory invasion may ease management through targeted interventions (Blackburn et al., 2011; Mack et al., n.d.).

The Shola Sky Islands (SSI) mountaintops of the Western Ghats (Robin & Nandini, 2012) - a UNESCO World Heritage site (UNESCO World Heritage Centre, n.d.), and global Biodiversity Hotspot (Myers et al., 2000) - are a good system to study understory invasion in long-established invasive woody stands. Centuries-old timber management has created a complex system of stands of invasive trees of three species (Arasumani et al., 2019; Joshi et al., 2018); *Acacia mearnsii*, one of the world’s most invasive species (*GISD*, n.d.), *Eucalyptus globulus*, and *Pinus radiata*, all known for their invasibility globally (Richardson & Rejmánek, 2011). They form widespread stands of single or double species in the SSI. Several other non-native species are known to colonise the understories of these stands with sporadic observations across the Shola Sky Islands ((Balaguru et al., 2016); (A. Das Pers. Obs.)). Species like *Ageratina spp*., *Ageratum spp*., *Cestrum aurantiacum, Lantana camara*, and *Pteridium aquilinum* (henceforth, Ageratina complex, Cestrum, Lantana, and Pteridium, respectively) have invaded multiple locations across SSI (Balaguru et al., 2016; Das, 2015). Several of these species are known to be noxious weeds globally (Cowie et al., 2018; Goncalves et al., 2014; Kaur et al., n.d.; Makokha, n.d.; Marrs & Watt, 2006; F. Wan et al., 2010). Most interactions between the invasive overstory species and non-native understory species are thought to be neutral or negative (Kuebbing & Nuñez, 2015), but positive interactions are not uncommon (Gómez-Aparicio, 2009). There is a need to highlight such associations to understand species ecology and manage the spread of invasives.

Based on the interactions between the over- and understory of exotic and invasive stands, the understorey community of colonising invasives varies between the types of stands (Power et al., n.d.; Wei et al., 2020). Attributes of non-native overstory trees that affect the understory community include canopy cover (Wei et al., 2020), nitrogen-fixing abilities (Kuebbing & Nuñez, 2015), basal area (Zilliox & Gosselin, 2014), soil-moisture (Wei et al., 2015), and allelopathy (Fisher, 1980). Sparser canopies in monocultures of Acacia, Pine and Eucalyptus, show greater regeneration capacity of heterospecific understory species (Arevalo et al., 2011; Forbes et al., 2016; Gwate et al., 2016). Acacia is capable of hindering the regeneration of heliophilic species (Gwate et al., 2016). In terms of allelopathy, Acacia stands can show extreme allelopathy, impeding heterospecific regeneration (Fatunbi et al., 2009). On the other hand, studies also indicate that these stands improve soil nitrogen dynamics (Forrester et al., 2007), which may improve regeneration conditions. Stands of *Pinus radiata*, show fewer allelopathic restrictions on the growth and germination of non-natives compared to *Acacia mearnsii* (Souto et al., 2001). *Eucalyptus globulus* stands show little evidence of allelopathy, and the soil in the stand usually promotes germination and seedling growth of other species (Nelson et al., 2021). Stands with individuals of Acacia and Eucalyptus trees may have better soil-based nitrogen than monocultures of *Eucalyptus globulus (Forrester et al*., *2007)*.

Invasive species show limited niche expansion beyond their native niches in environmental spaces (Liu et al., 2020). In novel or modified habitats of invasive stands, we expect the understory invasives to thrive in niche spaces similar to their places of origin. Differences in life forms of the colonising species may affect their response to underlying environmental gradients (Barbier et al., 2008; Zilliox & Gosselin, 2014) (refer to Supplementary Table 1). Here we propose to examine the spread of exotic invasive plant species in a mountainous, high-elevation landscape.

We identified non-native species colonising the understory of stands of exotic and invasive alien trees and conducted a region-wide assessment to answer the following questions:

1. Are there associations of secondary understory invasive species with specific invasive overstory species?
2. How do landscape factors and vegetation structure of exotic invasive stands affect the pattern of secondary invasion in the understory?

We expect plantation sites that provide similar microclimatic conditions associated with the original habitats of the colonising non-natives to demonstrate colonisation. With their sparser canopy cover and milder allelopathic effects, Eucalyptus stands may be expected to show higher species richness of non-natives and Acacia and Pine show the least species richness. Woody invasives may prefer warmer plots with greater human disturbances, while herbaceous invasives might thrive in open stands with higher soil moisture. Ferns may prefer wetter and warmer plots.

## Methods

We conducted this study across the high-elevation montane forests of the Nilgiris and Annamalai-Palani hills landscapes of the Shola Sky Islands (SSI) in the Western Ghats (10.12°N 77.60°E to 11.50°N 76.70°E) (inset A in Fig 1). We classified wooded habitats above 1400m MSL into shola forests (Robin & Nandini, 2012) and timber plantations, following (Arasumani et al., 2019).

**Fig 1:**
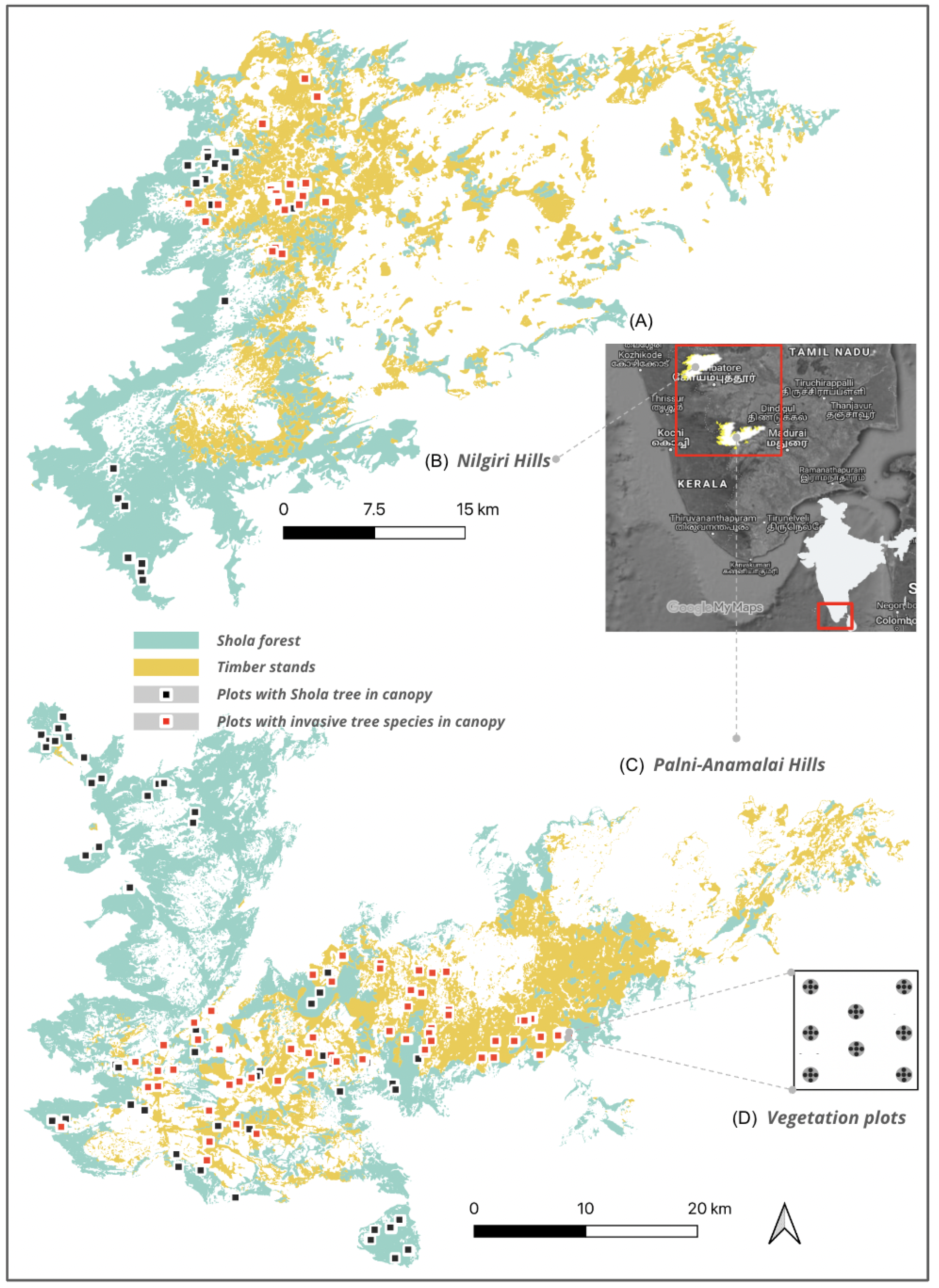
Location of study grids and vegetation plots. Inset map (A) shows the location of Shola Sky Islands of Nilgiri Hills (B) and Palani-Anamalai Hills (C), situated in southern India. The plots marked red have either Acacia, Pine or Eucalyptus as overstory trees. We sampled up to eight grids in each grid (D).

We divided the study region into 200m grids and randomly selected 0.5% of those (750 grids). We laid up to eight plots in 143 grids. We surveyed 232 plots with only exotic tree stands and no native tree species in the canopy. (Fig 1).

### Vegetation sampling

We sampled each grid with two to eight 7m-radius circular plots placed systematically (Fig 1 (D)). We identified and measured the circumference at breast height (GBH) for all live trees and snags over 30 cm within the circular plots (James & Shugart, 1970). In each plot, we placed 5 points (four at the edge along each cardinal direction and one at the centre). At these points, we measured the vegetation profile using a 5 m pole, calibrated at every 0.5 m (Karr, 1971). We noted calibrations at which vegetation touched the pole. The intensity of regeneration may depend on the amount of light reaching the floor, determined by the canopy cover, measured at those five points using HabitApp (Bianchi et al., 2017). At each of these points, we also noted the presence of recent fire incidents (burnt understory and blackened bases of the tree trunks) since fire is known to promote the spread of certain invasives, e.g. Pteridium (Carvalho et al., 2022) and control the spread of others, e.g. Lantana (Hiremath & Sundaram, 2005). We counted invasive species under the stands within a 1m radius plot around the five points: *Ageratum conyzoides, Ageratum houstonianum* and *Ageratina adenophora, Lantana camara, Cestrum spp*., *Solanum spp*., and *Pteridium aquilinum* and the conspecifics, *Acacia mearnsii, Eucalyptus globulus*, and *Pinus spp* and their heights were documented. Apart from these, other non-natives were also noted, and their heights were measured. We summed the number of leaf contacts in all strata within each plot along with the species-wise basal area and tree count. The canopy cover was averaged within each plot. We also summed the number of regeneration for each invasive species within each plot.

### Environmental variables

We selected putative predictors (described further below) for determinants of the intensity of regeneration (i.e., seed germination, seedling establishment and sapling growth) in plantations based on available literature. We assessed the following variables for each plot: elevation (Arévalo et al., 2005), slope (Cerdà & García-Fayos, 1997), aspect (Winkler et al., 2016), topographic position index (TPI) (Frey & Ashton, 2018), terrain ruggedness index (TRI) (Rhodes & St. Clair, 2018), topographic wetness index (TWI) (Petroselli et al., 2013) and topographic convergence index (TCI) (Bunn et al., 2005). We extracted topographic variables using the BHUVAN Cartosat-1 DEM v.3 tiles from tiles (30m contour interval) of our study area in QGIS3.10 Geospatial Data Abstraction Library (GDAL) plugin. We collected TPI, TRI and TWI at the local scale and TCI at scales of 30m, 60m and 90m after resampling the DEM. The aspect in radians was converted into two variables: the sine aspect (“eastness” of the plot) and the cosine aspect (the “northness” of the plot) because southern and western aspects in the Northern hemisphere receive greater solar irradiation (Piedallu & Gégout, 2008; Stage & Salas, 2007).

We calculated the area of shola forest within a buffer of 5ha around each plot (ln_shlbfr5ha) using data from (Arasumani et al., 2019). Within the same buffers, we extracted the lengths of roads for each plot rds_lngth_5ha. The data of the road network was procured from OpenStreetMap (https://www.openstreetmap.org) as a shapefile and clipped to the extent of the 5Ha buffers. The length of road networks within buffers may provide an indicator of the intensity of human access/disturbance for each plot (Benítez-López et al., 2010; Pauchard et al., 2009).

We also collected climatic variables for each plot studied from the CHELSA (Climatologies at High resolution for the Earth’s Land Surface Areas) data which consists of downscaled model outputs of temperature and precipitation estimates at a horizontal resolution of 30 arc sec (Karger et al., 2017). We extracted climatic variables influencing germination and plant growth: the temperature seasonality ((Wright, 1996)), precipitation in the dry quarters (e.g. (Howe, 1990; Martínez-Ramos et al., 2009)), precipitation in the cold quarters (Marques & Oliveira, 2008), minimum temperature in the cold quarter (Joshi et al., 2020), and the maximum temperature in the hot quarters (Wright, 1991).

### Analyses

To assess the adequacy of our sampling, we plotted the species-area curve for each stand type.

#### Ordination to represent the pairwise dissimilarity between sites

We assessed associations of understory non-native species with specific invasive overstory species using Non-metric Multidimensional Scaling (NMDS) with the Bray-Curtis distance as the dissimilarity measure (Minchin, 1987). Bray-Curtis is good at detecting underlying ecological gradients (Gauch, 1973). We used the package *vegan* (Oksanen et al., n.d.) with R (Team, 2013) and calculated the ordination using the function metaMDS. The dimensions were kept at 4, the maximum number of tries was 500, and the maximum number of iterations in the single NMDS run was 999. We visualised our NMDS plots using *ggplot2* (Wickham, 2016). We computed the subset of points on the convex hull created based on the categorical variable - the stand type. To statistically validate the composition affinity to the stand types, we performed the one-way ANOSIM nonparametric test with function *anosim* using *vegan (Oksanen et al*., *n*.*d*.). We examined two parameters for statistical significance and the degree to which the ordinations are related - a p-value and an R-value, respectively. R-values between 0.25 to 1 indicate a considerable difference (Polanía et al., 2020).

#### Phi Coefficient of association

The Phi coefficient of association (Tichy & Chytry, 2006) treats the target unit and the species symmetrically (joint fidelity measure), which means that an invasive taxon found exclusively in a particular stand type will have higher fidelity values. The strength of association between species assemblages and stand groups may indicate degrees of preferences for specific site groups. We used the function *‘multipatt’* of the package *indicspecies (De Caceres et al., 2016)*. We computed 95% confidence intervals using 999 permutations with the argument *func* as *r*.*g*.

#### Data Pre-treatment

We pooled the species *Ageratum conyzoides, A. houstonianum* and *Ageratina adenophora* (Ageratina complex), given very few records of the latter two. Our final list of species analysed included the following taxa: Ageratina complex, *Cestrum aurantiacum, Pteridium aquilinum*, and *Lantana camara*. All species in the overstory were removed from the data set: *Eucalyptus spp*., *Acacia mearnsii, Pinus spp*. We could not conduct analyses for *Solanum maritianum, Solanum sp*., *Ipomea sp, Clitoria ternatea, Tridax procumbens, Asparagus racemosum, Desmodium uncinatum, Oxalis corniculata, Urena lobata, Achyranthes aspera, Asteraceae, Urticaceae, Meliaceae, Malvaceae and Apiaceae* as the number of plots with these species regenerating was less than five. We removed the single plot with a mix of Eucalyptus-Acacia-Pine canopy from the analysis for Lantana alone.

Pearson correlation coefficient (r) was calculated among all pairs of variables. Using function *vif* from R package *usdm (Naimi et al*., *2014)*, we assessed multicollinearity further by calculating variance inflation factors for each invasive taxon (Dormann et al., 2013). The selected variables had pairwise r values lower than 0.7. The variance inflation factors were under 3.0 for the Ageratina complex, Cestrum, and Pteridium and under 7.0 for Lantana (Johnston et al., 2018). All chosen variables for all taxon are presented in Table 1.

Spatial autocorrelation for the regeneration data was assessed using the Mantel test with 1000 Monte Carlo permutations using *mantel*.*rtest* function from package *ade4 (Dray & Siberchicot, n*.*d*.). For each taxon, the simulated p-values were between 0.1-0.9, indicating no correlation between regeneration data (excluding zero regeneration) and the respective Euclidean distance matrices.

To account for the excess of absences of regeneration in the dataset, we fit a generalised linear mixed-effect model with zero-inflation to the data using the glmmTMB package (Magnusson et al., n.d.) in R. A plausible situation may be that some species do not regenerate in some regions because of environmental conditions or other factors, ultimately leading to the absence of that species in those regions. On the other hand, in some regions, even with facilitative conditions, the plots with zero regeneration can still occur by chance. The zero-inflation component models the absences in the former case while the count component models them in the latter case.

#### Model fitting

We tested the predictors for regeneration for each taxon in the count component of the model. For the zero-inflated component of the model, which models absence of regeneration, we chose TWI (proxy for soil moisture), road length within 5ha buffer (proxy for the source of propagules) and canopy cover (proxy for light intensity).

We rescaled continuous independent variables to zero mean and unit variance using the function *decostand* (vegan package). We used the glmmTMB package to fit the models and function *AIC* from package *stats* to extract each model’s Akaike Information Criterion corrected for small sample sizes (AICc). We fitted models in all possible combinations of the count component variables and the zero-inflation component variables for each taxon (Monteiro-Henriques & Fernandes, 2018), using the function *combn* from *utils* package (Maintainer, 2016).

#### Variable Importance Value (VIV)

We calculated the AIC differences to the best model for each fitted model. We extracted model weights with function *weights* in package *stat*s (Team et al., 2016). Variable importance values were computed at the end for each modelled variable by adding the Akaike weights of models consisting of the variable (Burnham et al., 2011; Monteiro-Henriques & Fernandes, 2018). We modified (Monteiro-Henriques & Fernandes, 2018) depiction of effect signals consistency with box plots for variable effects of each taxon studied. The signal of effect was considered consistent when the interquartile range did not overlap zero.

## Results

Of the 596 plots sampled, exotic timber trees formed the overstory in 232 plots, and these were selected for this study. Different exotic and invasive species individually and in combinations formed the overstory across this landscape - Eucalyptus 95 plots, Acacia 61, and Pine 28; Acacia + Eucalyptus 33 plots, other 7% of the total plots had other combinations of the three invasive species in the overstory.

### Which stands have the greatest number of secondary invasives?

We identified 24 species of invasives, at least to their family: *Ageratum conyzoides, A. houstonianum, Ageratina adenophora, Lantana camara, Cestrum aurantiacum, Solanum mauritianum, Solanum sp*., *Pteridium aquilinum, Ipomea sp, Acacia mearnsii, Eucalyptus globulus, Pinus spp, Clitoria ternatea, Tridax procumbens, Asparagus racemosum, Desmodium uncinatum, Oxalis corniculata, Urena lobata, Achyranthes aspera, Asteraceae, Urticaceae, Meliaceae, Malvaceae and Apiaceae*.

**Fig 2:**
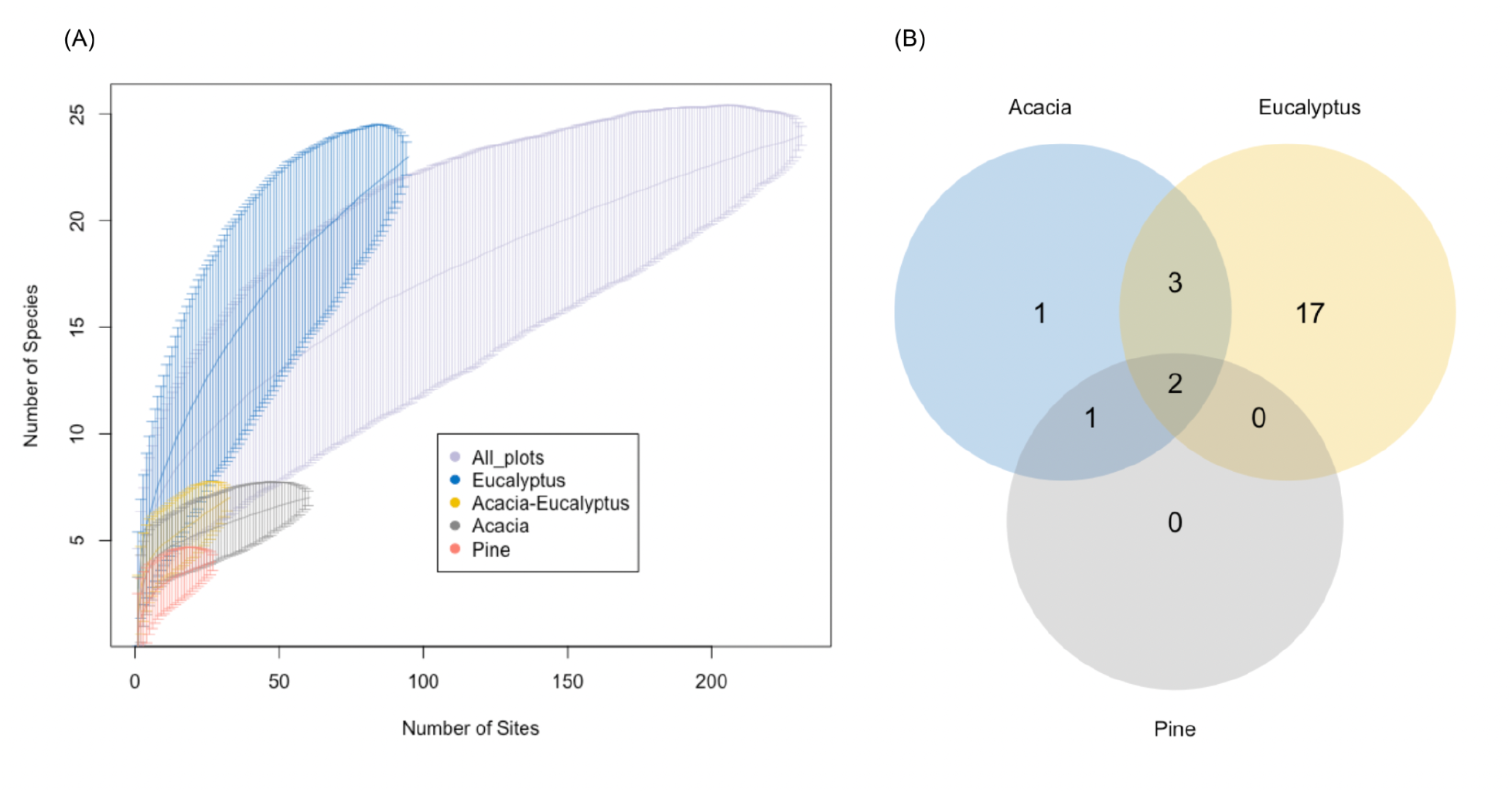
(A) Species accumulation curves for invasive regeneration in each exotic timber stand type, (B) Venn diagram representing the species richness of secondary invasives within each exotic overstory stand type

Eucalyptus stands host the maximum number of regenerating invasive species (17), followed by Acacia-Eucalyptus stands (3). Monocultural stands of Pine, did not have any invasive regeneration in the understory. The final NMDS stress was 0.05500354. Stress <0.1 provides good representation in reduced dimensions. The sites with different stand overstory types show a significant difference (ANOSIM; p<0.001, R=0.32) with regard to the invasive taxon regenerating in the understory (Fig 3: NMDS ordination).

**Fig 3:**
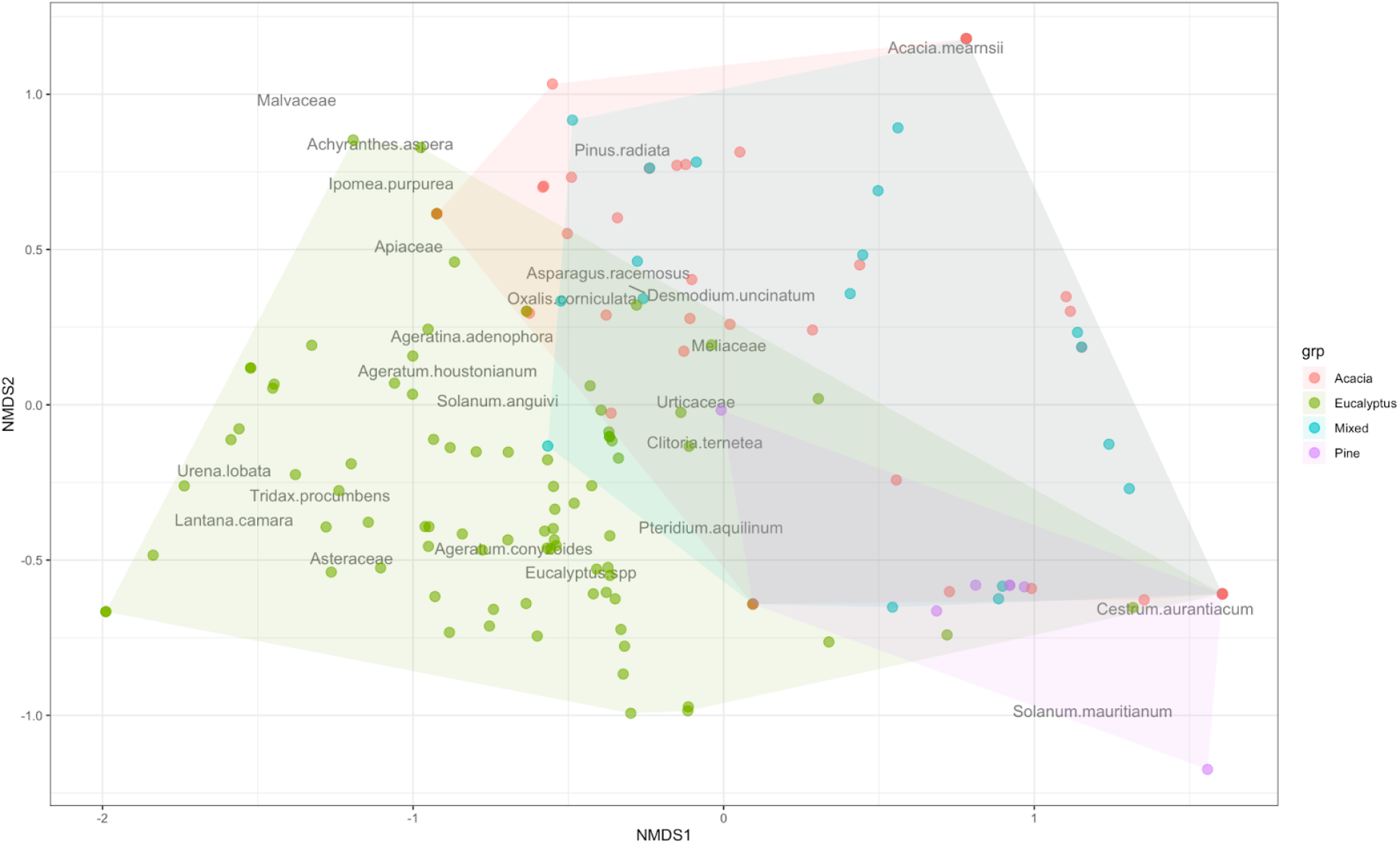
NMDS ordination: Invasive community analysed by stand type - Eucalyptus stands were different from Acacia and Pine which were more similar to each other. The closer points are more similar with respect to the presences and absences of the species within the plots.

The species-area curves were steep for stands of Eucalyptus and all stands considered cumulatively, but not for Acacia and Pine stands. This indicates that more invasive species may be hosted in the understory of exotic timber trees than we have uncovered with a sampling of 232 sites.

### Are there specific associations between exotic timber canopy species and understory invasives?

Species with fidelity values above 0.4 are usually diagnostic for the target vegetation unit (Stand types: Acacia, Eucalyptus, Pine, Mixed). *L. camara* was the sole taxon that showed high fidelity to Eucalyptus stands (fidelity value = 0.429, p=0.002) (Supplementary Table 3).

### Relationship of secondary invasives with habitat and environmental variables

Variable importance values based on Akaike weights identify the most relevant variables for the distribution of each invasive taxon in both - count and zero-inflation components (Supplementary Table 2).

*L. camara* was present in 41 plots with Eucalyptus canopy. Fire incidence and maximum temperature of the warm quarter were most important (Variable Importance Value = 82.8%, 78.4%, respectively); higher fire incidence was negatively correlated with the lantana occurrence, whereas higher dry quarter temperatures was negatively positively correlated with it. Canopy (VIV=99.9%) and road networks (VIV=97.9%) were important among the zero-inflation components. The effect signals were consistently positive for canopy cover but were inconsistent for the variable road networks (Figure 4). The former suggests a positive association of Lantana presence to canopy cover.

**Fig 4:**
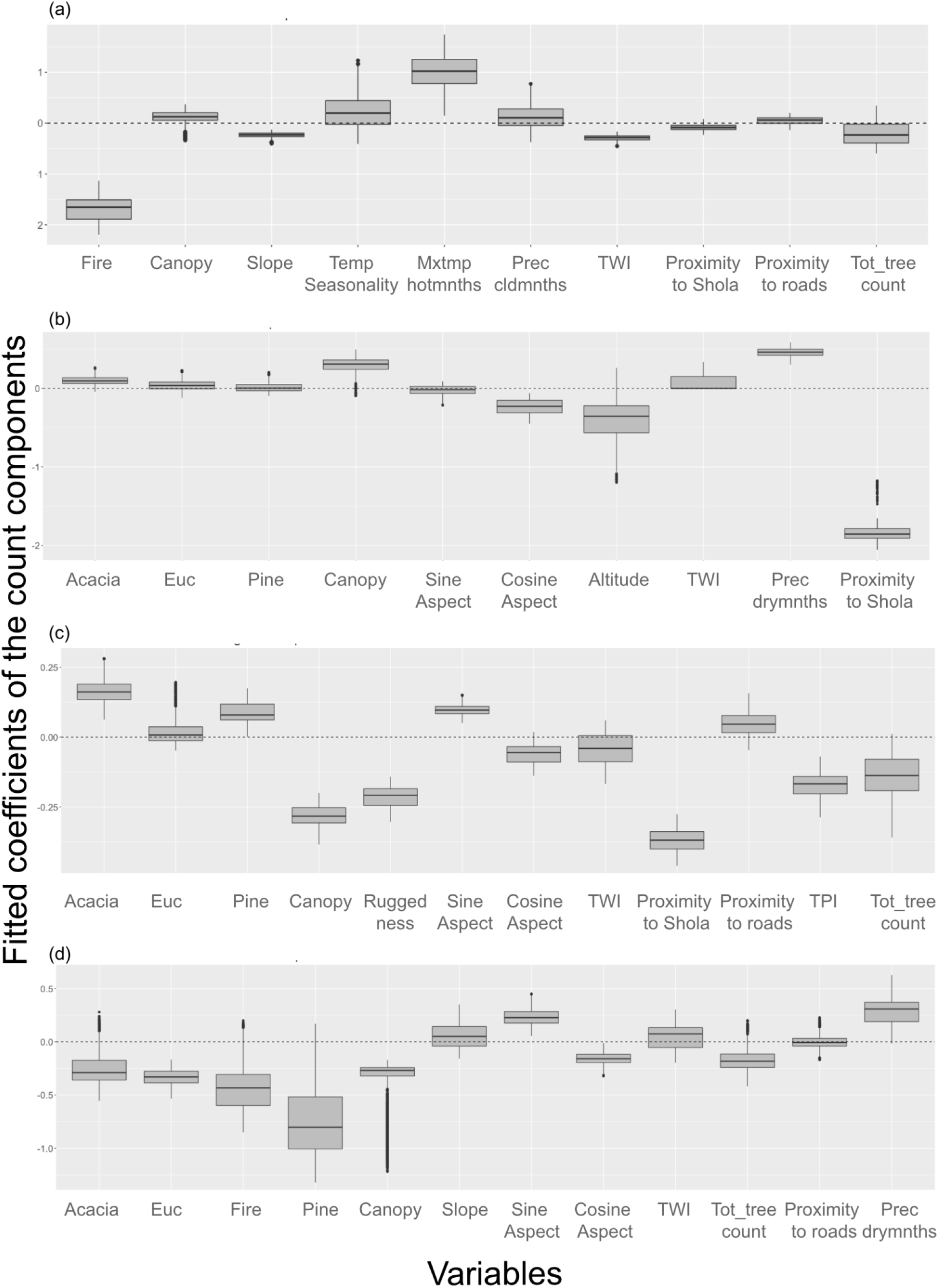
(a) Fitted coefficients of count components for Lantana sp., (b) Coefficients for Cestrum sp., (c) Coefficients for Ageratina complex, (d) Coefficients for Pteridium sp. With the importance value (Supplementary Table 2), we plotted the signal consistency of each variable: a box plot above the zero line indicates a consistent positive effect of the variable in the taxon regeneration; a box plot below the zero line indicates a consistent negative effect; if box plot contains zero, the signal was not consistent in the fitted models.

Cestrum was absent in our Palani-Anamalai hills plots and hence, this analysis is limited to the plots of the Nilgiri hill SSI. Cestrum was present in 45 plots out of the 79 plots used in analyses. Precipitation in the dry quarter, topographic wetness index, and the area of shola forest within a hectare around the study plots were important for the invasion of Cestrum (VIV= 88.1%, 87.8%, and 95.7%, respectively). Higher rainfall in drier months and greater soil moisture was positively associated with Cestrum occurrence, whereas proximity to Shola is negatively associated with it (Figure 4). Cestrum presence showed consistently negative effect signals for canopy cover (VIV for zero-inflation components 88.2%).

*Pteridium* was present in 92 plots. Basal area of Pine trees and fire are important determinants of fern invasion (count and zero-inflation components, respectively). *Pteridium* shows lower occurrence in plots with Pine presence, although the signal of the effects is inconsistent. Fire showed inconsistent signal effects for *Pteridium* presence.

*Ageratum conyzoides, A. houstonianum* and *Ageratina adenophora* in 83 out of 232 plots. The basal area of Acacia, canopy cover, proximity to shola, and terrain ruggedness index were important for the spread of Ageratina (79.3%, 99.5%, 99.9% and 84.0%, respectively). Plots with Acacia overstory had a higher probability of occurrence with regard to the Ageratina complex, and plots with greater canopy cover, proximity to shola forests and higher ruggedness had a lower probability of occurrence (Figure 4).

## Discussion

Our study examines the patterns of secondary invasion in the understory of plantations of exotic timber species, some of which are invasives themselves. We report a region-wide pattern of secondary invasion. Some of these patterns have important implications for ecological restoration actions that are currently underway in these habitats. All major exotic timber overstory species were associated with some secondary understory invasive species - Eucalyptus with Lantana, Acacia stands with the Ageratina complex, and Pine with Pteridium.

### Role of Eucalyptus

In our study, Eucalyptus stands show high species richness of other invasive species in their understory. This is a similar pattern to China, where understory invasion in the Eucalypt stands occurs significantly higher than in forests (Jin et al. 2015). In the Western Ghats, the spread of Lantana is a major threat (Joshi et al., 2015). Although Lantana has been known to invade lower elevations across a large part of India (Mungi et al., 2020), this study indicates the spread of Lantana to higher elevations as well, perhaps mediated by the presence of Eucalyptus akin to (O’Loughlin & Green, 2018). Apart from our study plots, we noticed Lantana in some of the higher reaches of the Western Ghats (∼1840 m ASL), although in very small patches indicating an emerging problem.

Some studies have noted a positive effect of Eucalyptus stands as they can have a nurse effect on the regeneration of natural forests (M. C. da Silva et al., 1995; Feyera et al., 2002). However, our results suggest that any natural regeneration of native species may be hampered by the invasives taking over these understories simultaneously.

#### Cestrum and other invasives

In our study, Cestrum was found only in the Nilgiri mountains, where the extent of invasion is alarming, ∼50% of all plots. It is widespread between 1800-2350m elevation (Das 2015). Although information is available indicating such invasion in the Nilgiris (Das, 2015; Mohandass & Davidar, 2009; Suresh et al., n.d.), a comparison of various montane areas was not conducted until now. Our findings indicate that unlike in the Nilgiris, Cestrum is not widespread in the Palnis, indicating regional differences in the spread of this invasive that requires further investigation. Canopy cover was negatively associated with the occurrence of both species - Lantana and Cestrum, and the Ageratina complex.

#### Implications for restoration

Shola Sky Islands have a large area (∼218 sq km) covered by non-native trees (Arasumani et al., 2019). Apart from the invasion into neighbouring shola grasslands of timber species like Acacia and Pine (Arasumani et al., 2019), our data indicate that these invasive timber plantations themselves appear to be hotbeds of invasion by other woody and herbaceous plants. The identity of these invasive species and patterns of invasion vary with the overstory species in the plantations.

The majority of our plots with Eucalyptus as the canopy species fall in the Western parts of the SSI system, which lies in the administrative boundary of the Kerala Forest Department. Apart from a different management system, these plots also have a heavier rainfall regime (Karger et al., 2017). Relative rainfall can have a positive association with lantana regeneration density (Debuse & Lewis, 2014). Increased light can result in increased cover of lantana in rainforests (Totland et al., 2005), but eucalyptus stands are relatively open in comparison with native forests. With low variability in the light intensity gradient, the lantana regeneration response may not be predominantly variable (Debuse & Lewis, 2014). In Pinus stands, leaf litter may hinder germination and hence colonisation of understorey species (Senbeta & Teketay, 2001). In older stands, the litter thickness is high and decomposition of such litter reduces with low temperature (Senbeta et al., 2002), possibly explaining the relative lack of invasive species in pine plantation understoreys. Finally, stands of *A. mearnsii*,, which are reported to show allelopathy, have a greater degree of shade, and lack of humidity in upper-soil (Tassin et al., 2009). Our study follows the above-mentioned patterns very closely.

At present, large areas are being planned for active restoration - removal of exotic, invasive timber trees - in both Kerala and Tamil Nadu (Correspondent, 2015). However, most action is targeted at Acacia and marginally at Pine. Here we present data showing that Eucalyptus plantations/stands should also be targeted for restoration when such activities are planned.

Eucalyptus species have been popular in compensatory afforestation programs across India (Vohra, 2021). Studies and reports abound in favour (Agarwal, S., & Saxena, A. K., 2017; *Website*, n.d.) and against the planting of Eucalyptus (Sikka et al., 2003). Our data suggest that, at least in some areas, these plantations can facilitate secondary invasion and should thus either not be permitted or actively managed to prevent invasion in the understorey.

We do note that our study was conducted only above 1400m elevation in the Western Ghats, and these patterns may be different elsewhere. We also note that this study does not present an exhaustive survey of invasives (see Supplementary Table 3 for details) or patterns of invasion in the region, but rather investigates specific ecological contexts of secondary invasives in an overstory of exotic and invasive tree plantations.

However, this study does indicate that large areas of the montane Shola Sky Islands that have been converted to exotic tree plantations are being impacted by secondary invasion of several species in ways that are specific to the habitat context and overstorey composition of these plantations.

## ACKNOWLEDGEMENTS

We thank the Forest Departments of Kerala and Tamil Nadu, specifically the Chief Wildlife Wardens, Ganga Singh IFS (Kerala) and Dr V. Naganathan (Tamil Nadu).

Our work would not be possible without the help from several specific members of Forests Departments - Kerala and Tamil Nadu, particularly Guru Swamy Dabbala (Nilgiri South DFO), V Ajayaghosh (Silent Valley assistant wildlife warden), R Lakshmi (Munnar Wildlife Warden), S.N. Thejaswi IFS (Tiruppur DFO), KK Kaushal IFS (FD Anamalai Tiger Reserve) and Siva Kumar (Ranger Kodaikanal). Our work was funded by the Ministry of Environment & Forests and Nat Geo Shola bird Ecology. Financial support from the DBT-RA Programme in Biotechnology and Life Sciences for AD is gratefully acknowledged. We received help for logistics of our field sampling from Vishnudas CK, Aravind PS, Tarsh Thekkekara, Vasanth Bosco, Ashwin Warudkar and Chiti Arvind and we are very grateful to them. Vasanth Bosco aso helped us in Shola species identification. We would also like to thank Devcharan Jathanna for his initial discussion regarding the study methodology.

This work spanned two states and involved several interns and field assistants as follows: Subash M, Karunan, Eswaran, Seetharaman, Kovai Thambi, Jegadeesh, Kamrajan, Preet Sharma, Anas Ibinu, Prithivraj B, Avinesh CN, Arjit Jere, Jude D, Roshan Robert, Mohammed Mubeen Mustafa, Arjun M, Vivek K Patel, Jay Prajapati, Krishna Priya M, Meghana Joseph, Nupur Kale, Ahmed Omar Haroon, Prerana Chandak, Kunal Gokhale, Rahul Mark N, Ritika Goswami, Shubham Suthar, Manav S, Abhishaek, Mikhail Nazareth, Sonia KB, Arathi J, Joseph Roy, Ahammed Saeed EP, Maithily Sawant, Keny J, Dhanesh.

## Notes

https://drive.google.com/file/d/1YRlwT0g8_s5bI2mfXDVmLqGmAB_Ip9xz/view?usp=share_link

### Competing Interest Statement

The authors have declared no competing interest.

