## Supplemental Tables 1, 2, 3 for "Patterns of secondary invasion in the understory of exotic, invasive timber stands"

### Supplementary

**Table 1: List of putative explanatory variables of Invasive regeneration and their categorization.**

| Model Component | Type of Variable | *Cestrum aurantiacum* | *Lantana camara* | *Pteridium aquilinum* | *Ageratina Complex* |
|---|---|---|---|---|---|
| **Count components** | **Compositional factors** | Acacia (Basal area) | | Acacia (Basal area) | Acacia (Basal area) |
| | | Eucalyptus (Basal area) | Eucalyptus (Basal area) | Eucalyptus (Basal area) | Eucalyptus (Basal area) |
| | | Pine (Basal area) | | Pine (Basal area) | Pine (Basal area) |
| | | Shola (Basal area) | | | |
| | **Structural factors** | Canopy cover | Canopy cover | Canopy cover | Canopy |
| | | | Total count of trees | Total count of trees | Total count of trees |
| | **Topographic factors** | Sine aspect | Sine aspect | Sine aspect | Sine aspect |
| | | Cosine aspect | Cosine aspect | Cosine aspect | Cosine aspect |
| | | Ln(elevation) | | | |
| | | | Slope | Slope | |
| | | TWI | TWI | TWI | TWI |
| | | | | | TRI |
| | | | | | TPI |
| | **Climatic factors** | | Temp Seasonality | | |
| | | | Precipitation in cold quarter | | |
| | | | Maximum temperature in hot quarter | | |
| | | | | Min temperature in cold quarter | |
| | | Precipitation in dry quarter | | Precipitation in dry quarter | |
| | **Landscape factors** | ln_shlbfr5ha | ln_shlbfr5ha | | ln_shlbfr5ha |
| | | | rds_lngth_5ha | rds_lngth_5ha | rds_lngth_5ha |
| | **Others** | | Fire | Fire | |
| **Zero-inflation component** | | Canopy | Canopy | | Canopy |
| | | TWI | TWI | | TWI |
| | | | rds_lngth_5ha | rds_lngth_5ha | rds_lngth_5ha |

| | | | | | |
|---|---|---|---|---|---|
| | | | | Fire | |
| **Random variables** | | Site | Type of plantation | Type of plantation | Type of plantation |
| **References for selection of variables** | | (Junaedi, 2013; Ojunga et al., 2020) | (Prasad, 2012; Sundaram & Hiremath, 2012) | (Amouzgar et al., 2020; Ú. S. R. da de Silva & Matos, 2006; Dolling, 1999) | (Lamsal et al., 2019; Parthasarathy et al., 2012; F. Wan et al., 2010; Yuan & Wen, 2018) |

**Table 2: List of Variable importance Values for Invasive regeneration**

| **Variable Importance Value** | | | | | |
|---|---|---|---|---|---|
| | **Variables** | **Lantana** | **Cestrum** | **Ageratina complex** | **Pteridium** |
| **Count component** | Acacia | NA | 28.0 | ***79.3*** | 44.9 |
| | Eucalyptus | NA | 22.4 | 26.5 | 68.1 |
| | Fire | ***100.00*** | NA | NA | 63.1 |
| | Pine | NA | 22.4 | 52.1 | ***75.7*** |
| | Canopy cover | 28.16 | 41.7 | ***99.5*** | 67.7 |
| | Slope | 39.46 | NA | NA | 29.6 |
| | Ln_elevation | NA | 29.4 | NA | NA |
| | Sine.aspect | NA | 21.9 | 44.5 | 53.3 |
| | Cosine.aspect | NA | 36.9 | 28.7 | 40.8 |
| | Temp.seasonality | 35.00 | NA | NA | NA |
| | Mxtmp_hotmnths | ***72.23*** | NA | NA | NA |
| | Prec_cldmnths | 28.49 | NA | NA | NA |
| | Prec_drymnths | NA | ***88.1*** | NA | 41.0 |
| | TWI | 47.14 | ***87.8*** | 27.5 | 29.0 |
| | Tot_ct_tree | 25.68 | NA | 33.1 | 32.0 |
| | ln_shlbfr5ha | 25.14 | ***95.7*** | ***99.9*** | NA |
| | rds_lngth_5ha | 28.89 | NA | 36.2 | 25.7 |

| | | | | | |
|---|---|---|---|---|---|
| | TRI | NA | NA | ***84.0*** | NA |
| | TPI | NA | NA | 45.9 | NA |
| **Zero-inflation component** | Canopy cover | ***99.95*** | ***88.2*** | 41.8 | NA |
| | TWI | 51.31 | NA | 40.2 | NA |
| | rds_lngth_5ha | ***98.39*** | NA | 51.4 | 37.2 |
| | Fire | NA | NA | NA | ***86.4*** |
| | ln_shlbfr5ha | NA | 32.8 | NA | NA |

**Table3: Association value between stand types and colonising invasive species, with heri statistical significance values. (Association value>0.4, usually diagnostic for the target vegetation unit )**

| Colonising species | Association value | Statistical significance |
|---|---|---|
| **Eucalyptus Stands** | | |
| *Lantana camara* | 0.486 | 0.001 |
| *Eucalyptus spp* | 0.459 | 0.001 |
| *Urena lobata* | 0.198 | 0.080 |
| *Ageratum conyzoides* | 0.191 | 0.094 |
| *Solanum sp* | 0.174 | 0.174 |
| *Ageratum houstonianum* | 0.148 | 0.321 |
| *Ipomea purpurea* | 0.116 | 0.594 |
| *Achyranthes aspera* | 0.111 | 0.611 |
| Malvaceae | 0.092 | 1.000 |
| Meliaceae | 0.092 | 1.000 |
| *Clitoria ternetea* | 0.092 | 1.000 |
| *Desmodium uncinatum* | 0.092 | 1.000 |
| *Asparagus racemosus* | 0.092 | 1.000 |
| *Oxalis corniculata* | 0.092 | 1.000 |
| Asteraceae | 0.092 | 1.000 |
| Apiaceae | 0.092 | 1.000 |

| | | |
|---|---|---|
| *Tridax procumbens* | 0.092 | 1.000 |
| Urticaceae | 0.092 | 1.000 |
| **Pine Stands** | | |
| *Solanum mauritianum* | 0.131 | 0.52 |
| *Pinus radiata* | 0.124 | 0.643 |
| **Acacia+Eucalyptus stands & Mixed stands** | | |
| *Acacia mearnsii* | 0.378 | 0.001 |
| **Eucalyptus & Pine Stands** | | |
| *Pteridium aquilinum* | 0.205 | 0.061 |
| **Acacia & Acacia+Eucalyptus & Eucalyptus stands** | | |
| *Ageratina adenophora* | 0.333 | 0.002 |
| **Acacia & Acacia+Eucalyptus & Pine Stands** | | |
| *Cestrum aurantiacum* | 0.263 | 0.015 |
